# Molecular mechanism of off-target effects in CRISPR-Cas9

**DOI:** 10.1101/421537

**Authors:** Clarisse G. Ricci, Janice S. Chen, Yinglong Miao, Martin Jinek, Jennifer A. Doudna, J. Andrew McCammon, Giulia Palermo

## Abstract

CRISPR-Cas9 is the state-of-the-art technology for editing and manipulating nucleic acids. However, the occurrence of off-target mutations can limit its applicability. Here, all-atom enhanced molecular dynamics (MD) simulations – using Gaussian accelerated MD (GaMD) – are used to decipher the mechanism of off-target binding at the molecular level. GaMD reveals that base pair mismatches in the target DNA at specific distal sites with respect to the Protospacer Adjacent Motif (PAM) induce an extended opening of the RNA:DNA heteroduplex, which leads to newly discovered interactions between the unwound nucleic acids and the protein counterpart. The conserved interactions between the target DNA strand and the L2 loop of the catalytic HNH domain constitute a “lock” effectively decreasing the conformational freedom of the HNH domain and its activation for cleavage. Remarkably, depending on their position at PAM distal sites, DNA mismatches leading to off-target cleavages are unable to “lock” the HNH domain, thereby identifying the ability to “lock” HNH as a key determinant. Consistently, off-target sequences hampering the catalysis have been shown to “trap” somehow the HNH domain in an inactive “conformational checkpoint” state (Dagdas et al. Sci Adv, 2017). As such, this mechanism identifies the molecular basis underlying off-target cleavages and contributes in clarifying a long-lasting open issue of the CRISPR-Cas9 function. It also poses the foundation for designing novel and more specific Cas9 variants, which could be obtained by magnifying the “locking” interactions between HNH and the target DNA in the presence of any incorrect off-target sequence, thus preventing undesired cleavages.

## Introduction

The CRISPR-(clustered regularly interspaced short palindromic repeats)-Cas9 is an adaptive immune system found in bacteria and archea conferring protection from foreign DNA.(1) By enabling deletion, insertion or correction of DNA at specific targeted sites within an organism’s genome, CRISPR-Cas9 is used as a genome editing technology holding enormous promises for medical, pharmaceutical and (bio)technological applications, while also being of invaluable impact for fundamental research.(1, 2) The CRISPR-Cas9 technology is based on a single protein – the endonuclease Cas9 – programmed with single guide RNAs (sgRNA) to site-specifically target any desired DNA sequence. The presence of a short sequence (i.e., a Protospacer Adjacent Motif, PAM) in close proximity to the cleavage site enables recognition of the desired DNA sequence across the genome and allows programmable applications. Upon PAM recognition, the DNA binds Cas9 by base pairing the RNA guide with one strand (the target strand, TS) and forming an RNA:DNA hybrid, while the non-target DNA strand (NTS) is unwound and also accommodated within the protein complex. Structures of the CRISPR-Cas9 complex indicate a recognition lobe, which mediates the binding of the nucleic acids through three REC1-3 regions, flanked by a PAM interacting (PI) domain and two nuclease domains, HNH and RuvC, which cleave the TS and NTS, respectively (Fig. 1A).

**Figure 1.**
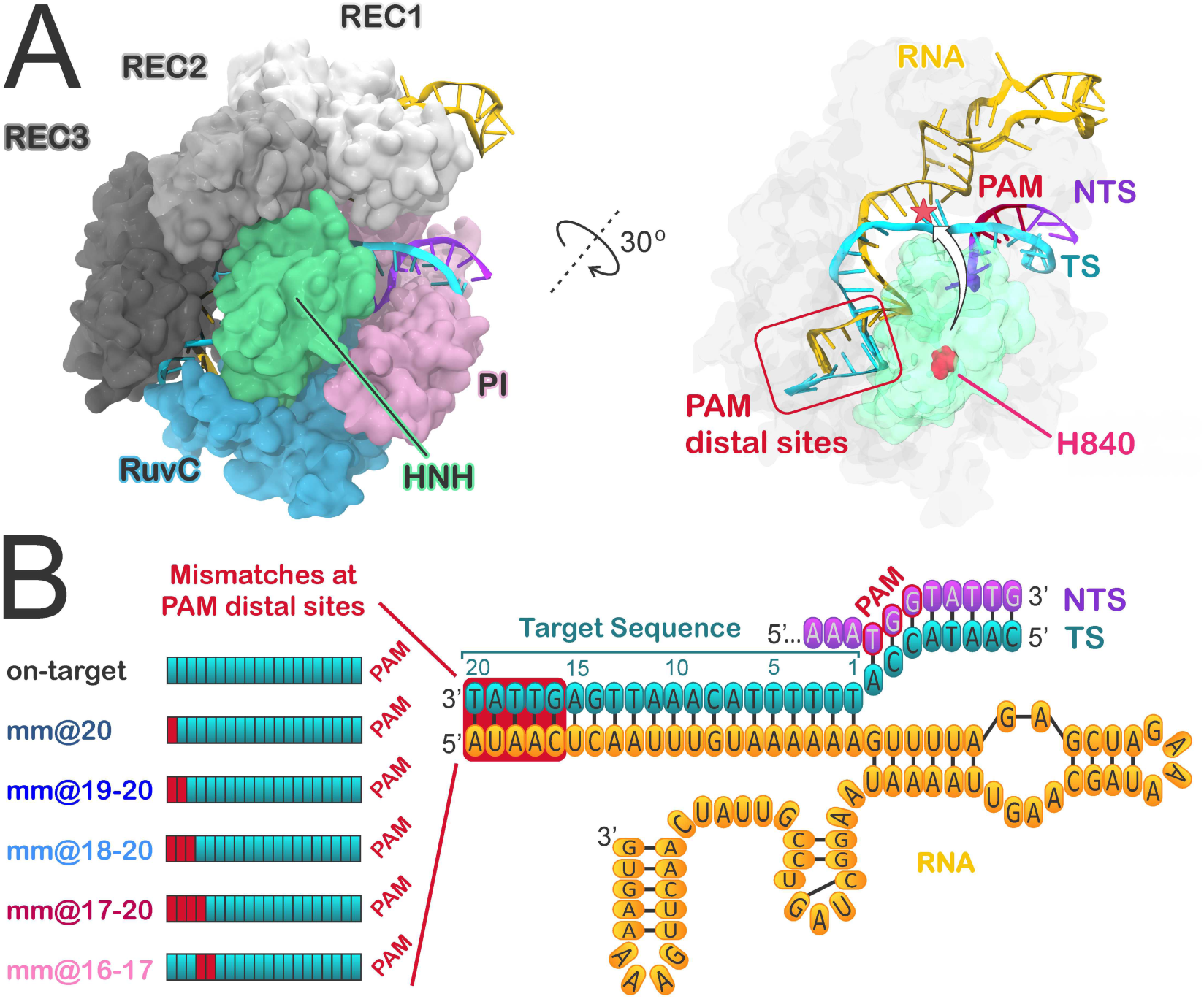
Overview of the CRISPR-Cas9 complex and of the simulated systems. **(A)** Crystal structure of the *S. pyogenes* CRISPR-Cas9 system, including the endonuclease Cas9, a guide RNA (orange), the target DNA (TS, cyan) and non-target DNA (NTS, violet) strands.(5) Cas9 is shown in molecular surface, with protein domains in different colors. The X-ray structure captures the inactive state of the HNH domain, which is a “conformational checkpoint” between DNA binding and cleavage. The right panel highlights the PAM distal sites on the RNA:DNA hybrid and the conformational change of the HNH domain required for catalysis (shown with an arrow), indicating the movement of catalytic H840 toward the cleavage site on the TS. **(B)** Diagram of the DNA and RNA filaments in CRISPR-Cas9, showing the location of base pair mismatches (mm) associated with off-target effects, at PAM distal sites. Six model systems were considered for MD simulations, including the on-target DNA sequence (on-target) and DNA sequences containing base pair mismatches at PAM distal sites (i.e., mm@20, mm@19-20, mm@18-20, mm@17-20 and mm@16-17).

In spite of the remarkable advantages of CRISPR-based genome editing with respect to traditional therapies, safety and efficacy issues have to be fully addressed prior to clinical applications of CRISPR-Cas9. The most severe issues limiting the applicability of CRISPR- Cas9 *in vivo* are the off-target DNA cleavages, which produce mutations at sites in the genome other than the desired target site, causing unwanted phenotypes.(3, 4)

In this respect, a promising strategy to fight off-target cleavages is the molecular engineering of highly specific Cas9 proteins,(6-8) which would enable safer and easy-to-use applications of the CRISPR technology on a genome-wide scale.(9) However, rational design of CRISPR-Cas9 requires detailed knowledge of the molecular bases underlying off-target effects, which are not yet understood.(10, 11) At the molecular level, off-target effects are the unselective cleavages of DNA sequences that do not fully match the guide RNA, bearing base pair mismatches at PAM distal sites of the DNA:RNA hybrid (Fig. 1). Kinetic and single molecule (sm) Förster Resonance Energy Transfer (FRET) experiments have shown that the occurrence of off-target cleavages is directly related to the conformational state adopted by the catalytic HNH domain.(7, 12) Upon DNA binding, HNH undergoes a conformational change from an inactive state, in which the catalytic H840 is far away from the cleavage site on the TS, to an activated state that is prone for catalysis (Fig. 1A). The inactive state of the HNH domain has been identified as a “conformational checkpoint” between DNA binding and cleavage, in which the RNA:DNA complementarity is recognized before the HNH domain assumes an activated conformation.(12) Sm FRET experiments have shown that the presence of DNA mismatches in the DNA:RNA hybrid can trap the HNH domain in the “conformational checkpoint” state, whose population increases by augmenting the number of mismatches at PAM distal sites. However, it is unknown how the presence of DNA mismatches can favor the inactivation of HNH. Detailed molecular knowledge of this mechanism is of major importance for developing more specific Cas9 variants, in which a single base pair mismatch is sufficient for trapping HNH in the inactive state, thus preventing the cleavage of any incorrect DNA sequence.

Here, we make use of extensive molecular dynamics (MD) simulations to characterize the molecular determinants of off-target binding, providing critical insights on the mechanism of off-target effects in CRISPR-Cas9. MD simulations have been shown to be a valuable tool for understanding the molecular basis of the CRISPR-Cas9 function.(13-17) Among other studies, our group has successfully applied MD simulations to disclose a mechanism for the conformational activation of Cas9, clarifying the activation process by which the HNH domain is repositioned for cleavage in good agreement with structural and smFRET experiments.(18) Remarkably, MD simulations as well as experimental studies have indicated that the conformational changes underlying the CRISPR-Cas9 function occur over and beyond micro-to-millisecond time scales.(13, 19, 20) Such long time scales require the use of computational methods that enable enhanced sampling of the configurational space, such as accelerated MD (aMD) simulations.(21) The aMD method adds a boost potential to certain regions of the potential energy surface, effectively decreasing the energy barriers and thus accelerating transitions between low-energy states. In this way, aMD simulations can capture biological processes occurring over milliseconds (and in some cases beyond), allowing the study of complex conformational transitions in folded or unstructured systems.(22) Recent advances have led to the development of a robust aMD methodology – i.e., Gaussian accelerated MD (GaMD)(23) – that extends the use of aMD to larger and more complex biological systems. Besides enabling the description of the activation mechanism in CRISPR- Cas9,(18) GaMD has been used to determine ligand binding in G-protein coupled receptors(24) and, remarkably, the mechanism of a G-protein mimetic nanobody binding of a medically important GPCR with intracellular signaling proteins, simulated for the first time.(25)

By using GaMD we determined the mechanism of binding of off-target sequences at the molecular level. GaMD simulations reveal how the presence of base pair mismatches at specific PAM distal sites of the RNA:DNA hybrid reduce the conformational mobility of the HNH domain and, in turn, its activation toward cleavage. We show that base pair mismatches at positions 17-16 promote newly formed interactions between the locally unwound TS and the L2 loop of the HNH domain, effectively decreasing its conformational mobility and preventing the HNH activation for cleavage. Being consistent with the available experimental data, the findings of this study outline a mechanism of off-target binding in CRISPR-Cas9, which poses the foundations for future structure based design of the system toward improved genome editing.

## Results

In order to determine how off-target sequences affect the conformational activation of the HNH domain, we performed all-atom GaMD simulations of the Cas9 protein in the “conformational checkpoint” state of the HNH domain (i.e., 4UN3.pdb).(5) GaMD simulations have already been used to successfully describe the activation process of the HNH domain(18) and are therefore ideal to characterize the effects caused by base pair mismatches in the conformational dynamics of CRISPR-Cas9 and its HNH domain. For this purpose, six model systems were built: one containing the fully matched RNA:DNA hybrid (considered the reference on-target system) and five systems containing base pair mismatches at different positions of the RNA:DNA hybrid (Fig. 1B). Specifically, we introduced 1 to 4 mismatches (mm) at PAM distal sites (i.e., at positions 20 to 16), resulting in the following models: mm@20, mm@19-20, mm@18-20, mm@17-20 and mm@16-17. These systems are consistent with experimental models for which the DNA cleavage rates have been measured.(7) Accordingly, the mm@20, mm@19-20 and mm@18-20 off-target systems cleave their DNA substrates with rates similar to the on-target system and are thus considered “productive” for DNA cleavage. Contrariwise, the mm@17-20 and mm@16-17 systems, which cleave their DNA substrates at significant slower rates, are considered “unproductive” for DNA cleavage. For each model system, >1 µs of GaMD was performed, with simulation conditions well suited for protein/nucleic acids complexes(26) and acceleration parameters that allow for sufficiently broad exploration of the conformational space and consistent observation of the molecular consequences of off-target DNA binding.

### DNA:RNA conformational dynamics in the presence of mismatches

During the dynamics, we observe an opening of the RNA:DNA hybrid at PAM distal sites with disruption of the Watson-Crick base pairing in the systems including off-target DNA sequences (Fig. 2). Contrariwise, in the system containing the on-target DNA, the DNA:RNA hybrid stably maintains its Watson-Crick base pairing, in good agreement with previous conventional and aMD simulations of CRISPR-Cas9.(13-18) Visual inspection of the trajectories reveals that all systems containing base pair mismatches, both “productive” and “unproductive” for DNA cleavage, display ill-behaved base pairs at the very end of the DNA-RNA hybrid (positions 20 to 18), whereas the on-target system remains well-behaved throughout the entire DNA:RNA hybrid (Fig. 2). The difference between “productive” and “unproductive” systems seems to lie at positions 17 to 16 of the RNA:DNA hybrid: at this region, only the “unproductive” systems (i.e., mm@17-20 and mm@16-17) display remarkable loss of the Watson-Crick base pairing (Fig. 2, right panels). At position 15, all systems keep their Watson-Crick base pairing intact, regardless of mismatches. It is important to note that transient openings at the end of a DNA duplex are not unusual in long timescale MD simulations, as shown by independent research groups including ours.(27-32) However, in the simulations of the on-target CRISPR-Cas9 system, the RNA:DNA hybrid maintains the Watson-Crick base pairing, most likely stabilized by the protein framework, as previously observed in conventional and aMD simulations of this system.(13-18) Clearly, the opening of the RNA:DNA hybrid at PAM distal sites only occurs in the presence of base pair mismatches.

**Figure 2.**
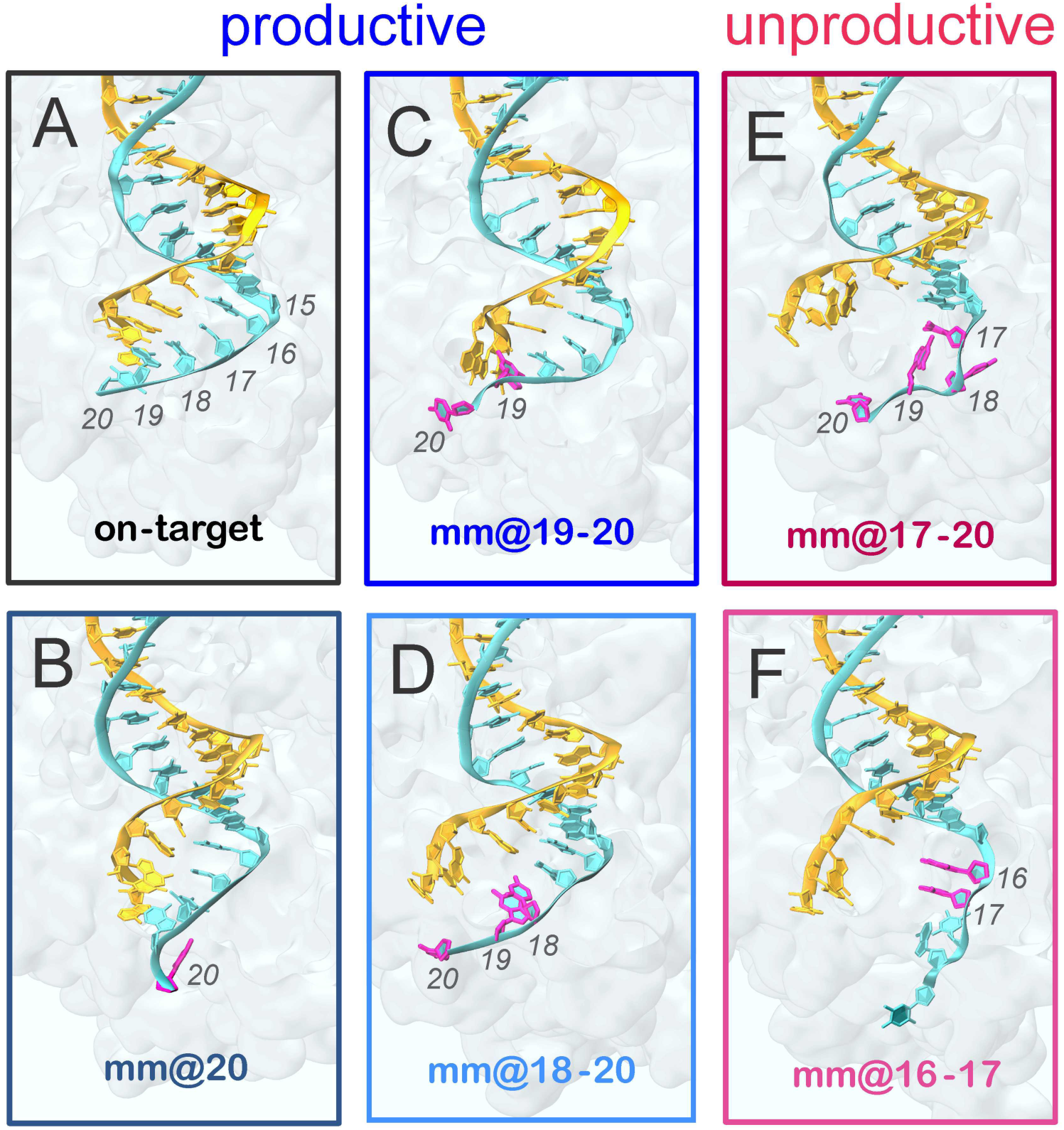
Conformations adopted by the RNA:DNA hybrid along GaMD simulations. Representative snapshots extracted from GaMD simulations of the CRISPR- Cas9 system, including the on-target DNA **(A)** and base pair mismatches (mm) at different positions of the hybrid: mm@20 **(B)**, mm@19-20 **(C)**, mm@18-20 **(D)**, mm@17-20 **(E)** and mm@16-17 **(F)**. “Productive” systems, which efficiently cleave their DNA substrate at rates similar to the on-target Cas9, are highlighted using cool colors, whereas “unproductive” systems, which slowly cleave their DNA substrate, are indicated with warm colors.(7) The RNA (orange) and the target DNA (TS, cyan) are shown as ribbons. Mismatched bases on the TS are highlighted in magenta. The protein environment is shown as a transparent molecular surface. These configurations are representative of the conformational changes detailed in Figs. S1-4 and in Fig. 3.

In order to better estimate the extent and the precise location of the nucleic acid distortions promoted by mismatches, we performed an in-depth analysis of the minor groove width at the PAM distal region of the DNA:RNA hybrid. In detail, the minor groove width has been computed at six different levels (*i-vi*), perpendicularly to the global helical axis (full details are in the Methods section). As an effect of the instability promoted by base pair mismatches, we detect an increase of minor groove width at PAM distal ends of the RNA:DNA hybrid in the systems containing off-target DNA sequences (Fig. S1). Interestingly, while “productive” systems promote the minor groove widening only at the very end of the hybrid (i.e., levels *i* and *ii*, Fig. S1), “unproductive” systems show a remarkable increase of the minor groove at more upstream regions of the hybrid, as shown by a shift of the probability distribution of the minor groove width toward larger values at the levels *iii*, *iv* and *v* (Fig. 3A). To complement the conformational analysis of the hybrid, the geometrical descriptors of base pair complementarity (Shear, Propeller, Stagger, Buckle, Stretch, Opening) have been computed. At position 17 (Fig. 3B), the broadly distributed scatter plots produced by “unproductive” systems (i.e., mm@17-20 and mm@16-17) are indicative of significant loss of base pairing. Contrariwise, systems that are “productive” for DNA cleavage (on-target, mm@20, mm@19-20 and mm@18-20) display confined distributions of scattered dots, as consistent with well-matched base pairs. Figs. S2-S4 report data for all simulated systems computed at positions 20 to 15, revealing that the loss of complementarity at the very end of the hybrid (i.e., positions 20 to 18) is observed in all off-target sequences, while only the “unproductive” systems show remarkable loss of the complementarity at positions 17 to 16. At position 15 all systems keep their Watson-Crick base pairing intact, consistently with the visual inspection of the trajectory (Fig. 2).

**Figure 3.**
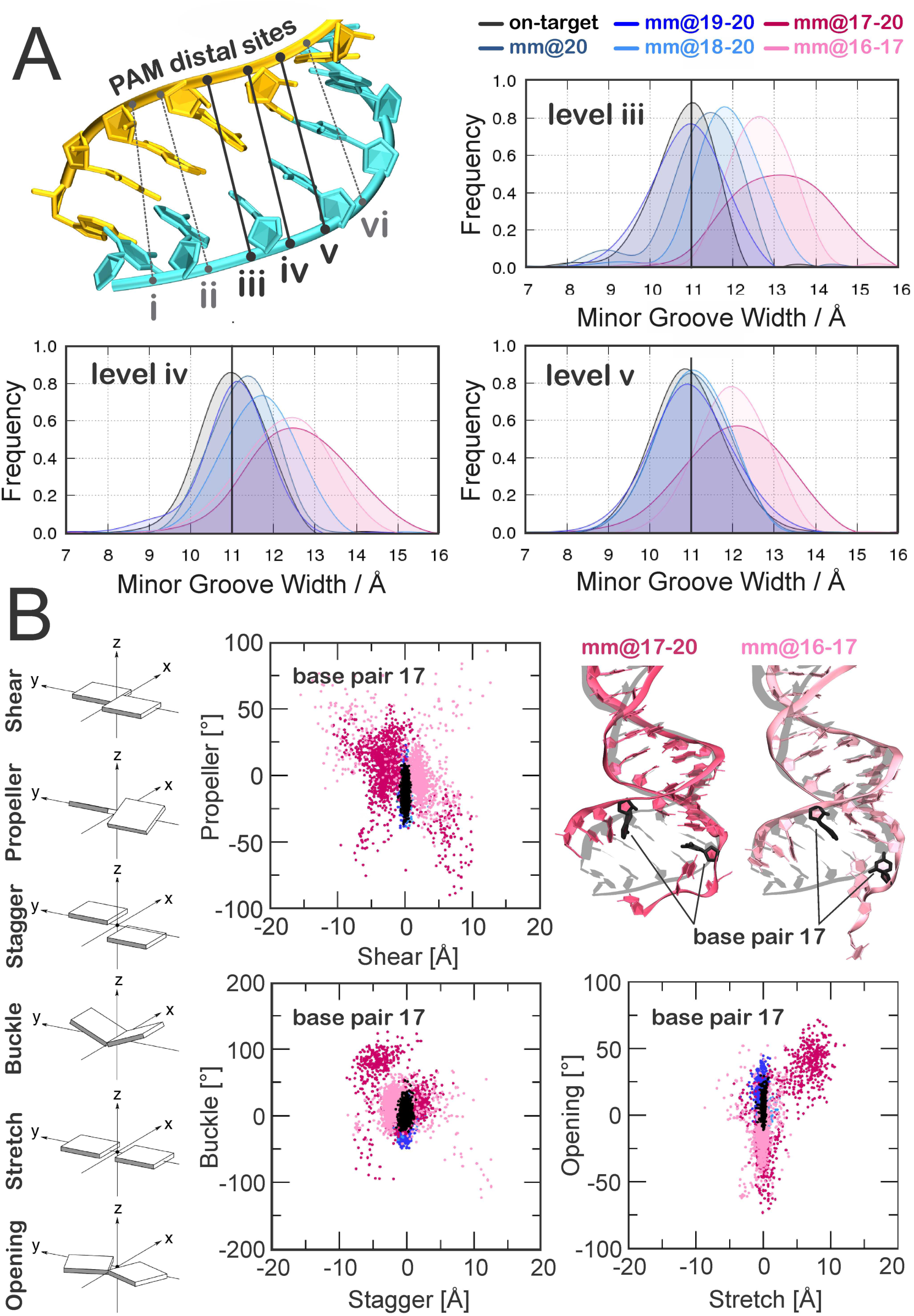
RNA:DNA geometrical properties along GaMD simulations. **(A)** The minor groove width in the PAM distal region of the RNA:DNA has been computed at six different levels (i-vi), which are schematically shown on the 3D structure of the RNA:DNA hybrid. The probability distributions of the RNA:DNA minor groove width at the (iii-v) levels are shown for the on-target CRISPR-Cas9 and for the systems including base pair mismatches at different positions of the hybrid (i.e., mm@20, mm@19-20, mm@18-20, mm@17-20 and mm@16-17). A vertical bar indicates the experimental minor groove width (i.e., ∼11 Å from x-ray crystallography, enlarged by ∼1 Å if NMR data are considered).(28) Full data are reported in Fig. S1. **(B)** Scatter plots of the geometrical base pair descriptors, computed at position 17 of the RNA:DNA hybrid for all studied systems (full data are in Figs. S2-4), in the same color scheme used in panel ATranslational (i.e., Shear, Stretch and Stagger) and angular (i.e., Buckle, Propeller and Opening) descriptors are expressed in Å and degrees, respectively. The RNA:DNA hybrid is shown on the right for the on-target CRISPR-Cas9 (gray) superposed to the mm@17-20 (red) and mm@16-17 (pink) systems.

Taken together, the base pair geometrical descriptors well agree with the analysis of the minor groove widths, indicating a significant minor groove widening promoted by “unproductive” mismatches at levels *iii*, *iv*, and *v* (Fig. 3A) and the loss of base pairing at position 17 (Fig 3B), which is not observed in the “productive” systems. Overall, as an effect of base pair mismatches at positions 16 to 20, “unproductive” systems show an extended opening and distortion up to position 17 of the RNA:DNA heteroduplex. These results pinpoint to a general trend distinguishing “productive” and “unproductive” mismatches, where the extent and precise location of the distortions in the hybrid plays a key role for DNA cleavage efficiency. Specifically, mismatches that perturb the hybrid downstream of position 17 are “productive” for DNA cleavage, while mismatches whose distortions occur (or are propagated) to position 17 are “unproductive” for DNA cleavage, as is the case of mismatches mm@17-20 and mm@16-17 (Fig. 2).(7)

### Effect of off-target binding on the catalytic domains

The observed opening of the RNA:DNA hybrid causes novel interactions with the protein framework. Here, we detail the interactions established by the RNA:DNA hybrid with the catalytic domains of Cas9, HNH and RuvC. Specifically, we measure the tight contacts (i.e., within 4 Å radius) established along the dynamics by the hybrid and the neighboring residues.(33) As a result, we detect a remarkable increase of interactions between the hybrid and the HNH domain in the “unproductive” systems (mm@17-20 and mm@16-17), which is particularly relevant with polar and charged residues (Figs. 4 and S5). Remarkably, the relevance of these newly formed contacts is confirmed by the statistical error non-overlapping with the “productive” systems. An increase of interactions for the “unproductive” systems is also observed with the RuvC domain. The interaction of the RNA:DNA hybrid with the HNH/RuvC domains in the mm@17-20 system is shown in Fig. 4 (right panel). Importantly, the HNH domain connects RuvC through two flexible loops: L1 (residues 765-780) and L2 (residues 902-918), which are part of HNH and are important in the function of CRISPR- Cas9 (Fig. 4A, right panel). Indeed, L1/L2 intervene in the repositioning of HNH from the inactivated (i.e., “conformational checkpoint”) to the activated state, by changing configuration.(18, 34) As well, L1/L2 exert an allosteric control on the activity of HNH and RuvC, enabling the information transfer for concerted cleavages of the two DNA strands.(15, 35) In order to understand the role of these two loops in the interaction with the hybrid, we specifically measured the interactions established by L1/L2 and the DNA:RNA hybrid. As a result, the interactions of L1 do not show relevant differences among the investigated systems (Fig. S5B). Moreover, L1 is a disordered loop that is highly flexible, resulting in overlapping error bars. Contrariwise, the interactions established by the RNA:DNA hybrid and L2 are remarkably increased in the “unproductive” systems (Fig. 4 and S5B). As noted above, the increase in interactions involving L2 in “unproductive” systems has non-overlapping error bars with the “productive” systems and is particularly relevant for polar and charged residues.

**Figure 4.**
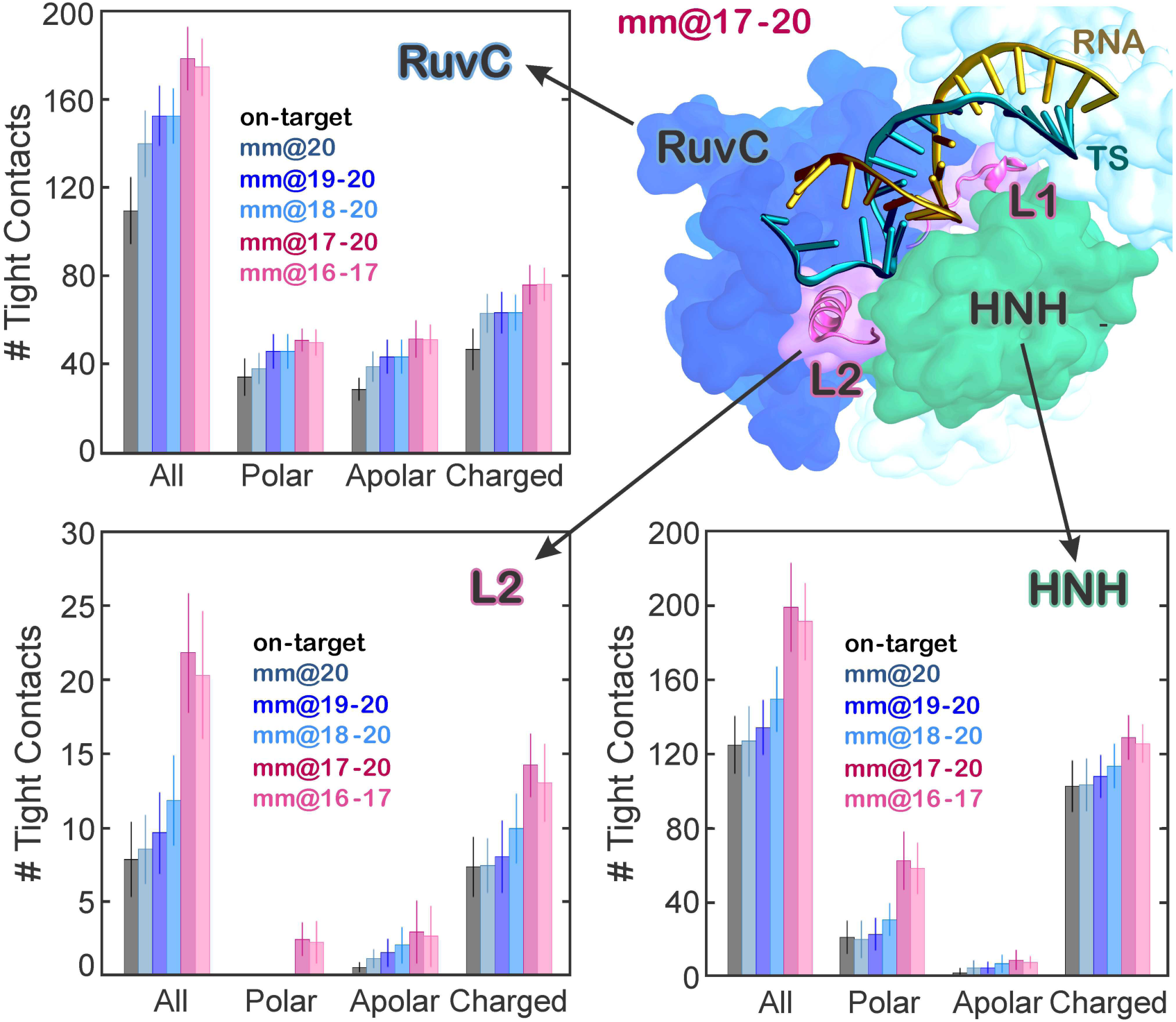
Quantitative evaluation of the interactions between the RNA:DNA hybrid and the Cas9 protein. Number of tight contacts (i.e., within 4 Å radius) established along the dynamics by the PAM distal ends of the RNA:DNA hybrid with the neighboring residues of the HNH and RuvC domains, as well as with the L2 loop connecting HNH to RuvC. The total number of interactions was also separated in polar, apolar and charged contributions. A cartoon of the mm@17-20 system, highlighting the RNA:DNA hybrid and its interactions with HNH and RuvC, as well as with the L1/L2 loops, is shown (right panel).

Analysis of the interactions between the hybrid and the HNH domain excluding the residues belonging to L2 confirms that the increase of interactions between HNH and the hybrid occurs at the level of L2 – practically vanishing when L2 is excluded from the analysis – and mainly involves polar and charged residues (Fig. S5B). A more detailed inspection of the trajectories reveals specific interactions that are exclusively formed in the “unproductive” systems and could be key for explaining their reported slow DNA cleavage rates.(7) We detect the formation of a salt bridge between Arg904 of L2 and the TS backbone at position 17 (Fig. 5, shown for the mm@17-20 system). Remarkably, this interaction with the TS backbone is formed along the simulations of both “unproductive” systems (mm@17-20 and mm@16-17), whereas it does not form in the on-target Cas9 or in the “productive” off-target systems (Fig. S6). In the mm@17-20 system, additional interactions are also established upon ∼0.45 µs between the Asp911 and Ser908 residues of L2 and the base 18 of the TS, which is flipped out of the hybrid (Fig. 5). Remarkably, the interaction with the TS backbone is conserved in the “unproductive” systems, as it is favored by the local unwinding of the DNA:RNA hybrid at position 17, which is in turn promoted exclusively by the “unproductive” mismatches (i.e., mm@17-20 and mm@16-17, Figs. 2 and 3). Contrariwise, when the hybrid is well formed at this specific level (i.e., position17), as in the case of the “productive” mismatches and for the on-target DNA, the TS backbone is not free to approach and bind the L2 loop.

**Figure 5.**
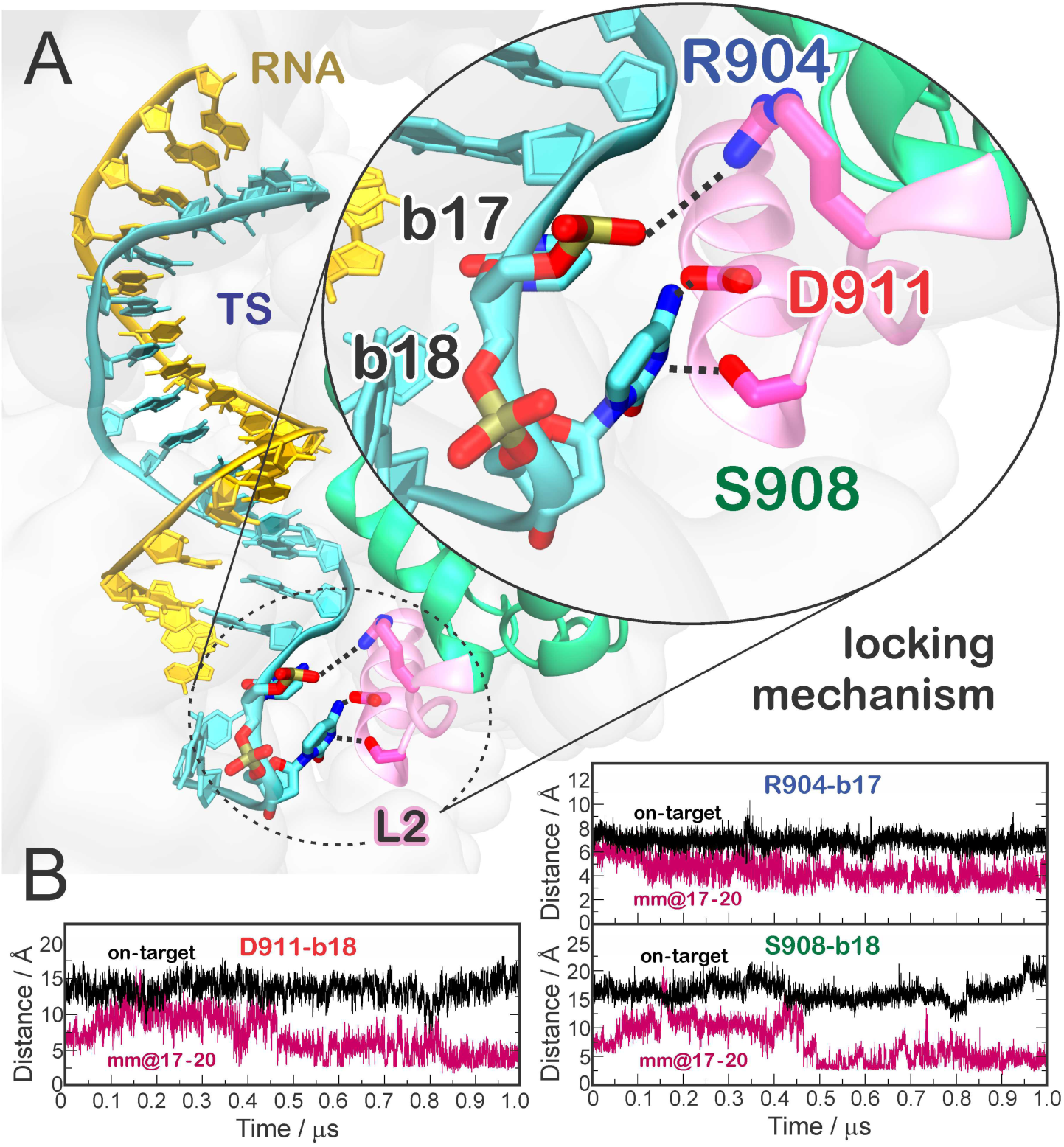
Locking interactions between the DNA target strand (TS) and the L2 loop of the HNH domain. **(A)** Representative snapshot of the mm@17-20 system, showing the interactions established by the RNA:DNA hybrid with residues of the L2 loop. **(B)** Time evolution of the interactions established by R904, D911 and S908 and the TS at positions 17 (R904) and 18 (D911 and S908). Data for the on-target Cas9 (black) are compared to the mm@17-20 system (red).

When formed, these interactions constitute a “lock” for the L2 loop, decreasing its conformational flexibility. This is confirmed by a lower root mean square deviation (RMSD) of the L2 loop in “unproductive” systems as compared to the on-target system (Fig. S7). Considering that the activation of the HNH domain requires a substantial conformational rearrangement of L2,(18, 34, 35) which moves away from the PAM distal region of the hybrid, the newly formed interactions observed in the “unproductive” systems can prevent HNH from undergoing its conformational activation. This mechanism is consistent with several experimental studies showing that the presence of base pair mismatches up to positions 17-16 trap HNH in a “conformational checkpoint” state, preventing it from reaching the active conformation.(7, 12, 36)

On the contrary, the presence of up to 3 mismatches at positions 20-18 still allows HNH to correctly reposition for cleavage, although with slightly slower rates as compared to on-target DNA sequences. Remarkably, this experimental evidence is consistent with the outcomes of GaMD, showing that 3 mismatches at positions 20-18 are unable to “lock” the HNH domain (Fig. S6). In light of these evidences, our simulations reveal the key interactions – formed between a locally unwound TS and the L2 loop – by which specific base pair mismatches at positions 17-16 prevent HNH activation for cleavage.

### Effect of off-target binding on the hybrid recognition

The formation of the RNA:DNA hybrid has been shown to be a prerequisite for the on-target selectivity.(7) Single molecule and bulk experiments have shown that the REC lobe of CRISPR-Cas9 exerts a key role in the recognition. Specifically, the REC3 region, which directly contacts the very end of the hybrid, “senses” the formation of the DNA:RNA hybrid and allows for HNH nuclease activation.(7, 13)

In order to understand the role of REC3 in the hybrid recognition and in the on-target selectivity, we monitored the interactions established by the RNA:DNA hybrid with REC3 in the simulated systems. In the “unproductive” systems, we observe a significant increase of interactions between the hybrid and REC3 (separated in polar, apolar and charged contributions, Fig. 6A). In these systems, due to the extended opening of the RNA:DNA hybrid, the 692-700 α-helix inserts within the heteroduplex, causing novel steric and electrostatic interactions (Fig. 6B). On the contrary, in the on-target Cas9, as well as in the “productive” systems including off-target sequences, the 692-700 α-helix does not insert within the hybrid (Fig. 6C). The 692-700 α-helix includes a set of key residues (N692, M694, Q695 and H698), whose mutation to alanine confers increased selectivittarget against off cleavages.(7) Indeed, the hyper accurate Cas9 (HypaCas9) variant, including the N692A, M694A, Q695A and H698A mutations, cleaves the on-target DNA with rates similar to the wt-Cas9, but the cleavage is reduced in the presence of mismatches. The specific interactions of these residues have been monitored during GaMD. As a result, in the on-target Cas9, N692 and Q695 stably bind the TS backbone thoughout the dynamics, while M694 and H698 establish additional interactions (Fig. S8). Remarkably, the interactions established by N692 and Q695 with the TS are also observed in the “produtive” off-target systems, but are lost in the “unproductive” systems (Fig. S9).

**Figure 6.**
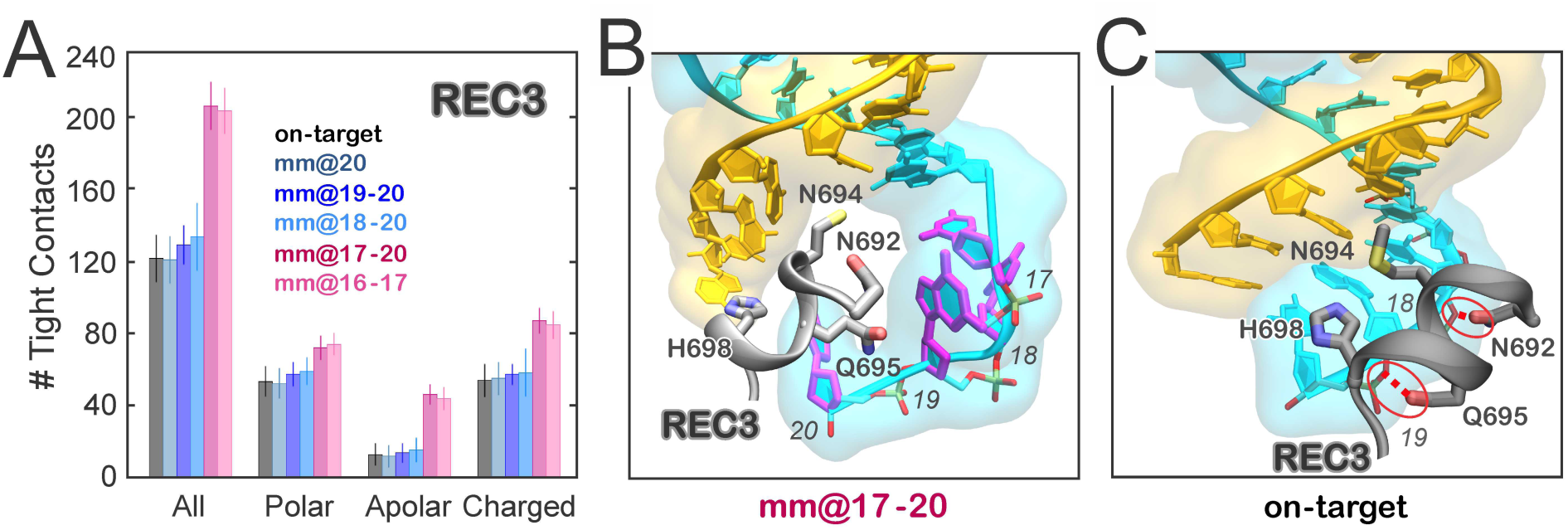
Role of REC3 in the off-target binding. **(A)** Number of tight contacts (i.e., within 4 Å radius) established along the dynamics by the RNA:DNA hybrid and the neighboring residues of the REC3 region, computed considering all residues, as well as the groups of polar, apolar and charged residues. **(B)** Representative snapshot from GaMD simulations of the mm@17-20 system, showing the extended opening of the RNA:DNA hybrid and the insertion of the 692-700 α-helix (gray) within the heteroduplex. **(C)** Snapshot from GaMD of the on-target CRISPR-Cas9, showing a well-behaved RNA:DNA hybrid and the conserved interactions established by Q965, N692 and the DNA TS. The RNA (orange) and the DNA TS (cyan) are shown as ribbons. The 692-700 α-helix (gray) is shown as cartoon; interacting protein residues are shown as sticks.

These data indicate that the 692-700 α-helix is a key element for the recognition of the complementarity of the RNA:DNA hybrid and the selection between “productive” and “unproductive” systems. Indeed, in the “unproductive” systems, the insertion of the α-helix 692-700 within the heteroduplex contribues to the observed opening of the RNA:DNA hybrid and, in turn, to the establishment of interactions between the TS and the L2 loop (Fig. 5). This establishes a mechanism of selectivity where, in the “productive” systems REC3 “senses” a well-formed hybrid through the 692-700 α-helix. Indeed, in the on-target Cas9 and in the presence of 1 to 3 mismatches at PAM distal ends, the N692 and Q695 stably bind the TS backbone, contributing also to the stabilization of the hybrid itself. Contrarywise, in the presence of an extended opening of the RNA:DNA hybrid, as including base pair mismatches at positions 17-to-16, the 692-700 α-helix moves apart losing its interaction with the TS backbone (Fig. S9) and results in the insertion within the hybrid.

Overall, these observations provide a rationale for the role of REC3 in the on-target selectivity, revealing how by “sensing” the RNA:DNA hybrid, REC3 discriminates “productive” from “unproductive” sequences for cleavage. Finally, in light of these observations, it is likely that the mutation of the N692, M694, Q695 and H698 residues to alanine, as in HypaCas9,(7) would reduce the electrostatic interactions with the hybrid, altering the mechanism depicted above. Although likely, this speculation will require extensive GaMD simulations of HypaCas9, which could ultimately capture the mechanism of altered specificity.

## Discussion

### Molecular basis of off-target binding

Off-target effects represent a severe issue hindering a full exploitation of the CRISPR- Cas9 technology in *in viv*o applications.(3, 4, 6-8) Here, by using accelerated MD simulations we capture the mechanism of binding of off-target sequences at the molecular level. A Gaussian accelerated MD methodology(23) – enabling to capture long time scale conformational changes that are not accessible via conventional MD simulations – has been applied to characterize the binding of off-target DNA sequences, compared to a fully matching on-target DNA.

GaMD reveals that while the on-target DNA remains fully matched to its complementary RNA, the presence of base pair mismatches induces an opening of the RNA:DNA hybrid (Fig. 2). We observe that 1 to 3 mismatches up to position 18 perturb the hybrid at its very end, whereas base pair mismatches up to positions 17-16 produce an extended opening of the RNA:DNA heteroduplex. Remarkably, the presence of 1 to 3 base pair mismatches up to position 18 at PAM distal sites still enables productive cleavages of the DNA substrate at similar rates of the on-target Cas9.(7) Contrariwise, by increasing the number of base pair mismatches up to positions 17-16 or by introducing two mismatches specifically at positions 17-16, Cas9 is rendered unproductive, exhibiting slower DNA cleavage rates. This observation identifies a distinction between “productive” and “unproductive” mismatches, where the extent and precise location of the distortions in the hybrid plays a key role for DNA cleavage efficiency.(7) Indeed, mismatches that perturb the hybrid downstream of position 17 are “productive” for DNA cleavage, while mismatches whose distortions occur (or are propagated) up to positions 17 and 16 are “unproductive” for DNA cleavage.

As an effect of the extended opening observed in the presence of base pair mismatches at positions 17 and 16, the DNA TS engages in conserved interactions with the L2 loop of the HNH domain (Fig. 5). In particular, Arg904 of the L2 loop stably binds the TS backbone at position 17 in both the “unproductive systems” studied herein, while additional interactions with the TS bases also involve Asp911 and Ser908. These interactions constitute a “lock” for the L2 loop, decreasing its conformational freedom (Fig. S7). Considering that the activation of the HNH domain requires the conformational rearrangement of L2,(18, 34, 35) which enables HNH to properly relocate for the TS cleavage, the newly formed interactions prevent HNH from undergoing its conformational activation. This indicates that base pair mismatches at positions 17 and 16 of the RNA:DNA hybrid hinder the activation of HNH through the binding of L2. This mechanism is consistent with the experimental evidence that base pair mismatches up to positions 17-16 of the hybrid traps HNH in an inactive “conformational checkpoint” state, preventing it from reaching the active conformation.(7, 12, 36) Contrariwise, the presence of up to 3 base pair mismatches at positions 20 to 18 does not result into stable interactions “locking” L2 (Fig. S6), as revealed by GaMD simulations. This is in agreement with the fact that 3 mismatches (at positions 20-18) still allow the repositioning of HNH (although slower than in the presence of the on-target sequence) and are “productive” for DNA cleavage.(7, 12, 36) In light of these experimental evidences, GaMD simulations depict a mechanism for the binding of off-target sequences at PAM distal sites, capturing the interactions – formed between a locally unwound TS backbone and the L2 loop – by which base pair mismatches at the specific positions 17-16 prevent HNH activation for cleavage. GaMD also shows that the REC3 region of the Cas9 recognition lobe undergoes a significant conformational rearrangement in the presence of base pair mismatches up to position 17 and 16 of the RNA:DNA hybrid (Fig. 6). Indeed, the 692-700 α-helix of REC3 inserts within the heteroduplex, further contributing to the observed opening of the RNA:DNA hybrid and, in turn, to the establishment of interactions between the TS and the L2 loop. Contrariwise, in the presence of an on-target DNA, as well as sequences including up to 3 base pair mismatches at PAM distal sites, the 692-700 α-helix does not insert within the hybrid, while the N692 and Q695 residues stably bind the TS backbone. This clarifies the mechanism by which REC3 “senses” the formation of a formed hybrid through the 692-700 α-helix,(7, 13) contributing also to the stabilization of the hybrid itself.

Here, GaMD simulations have established a mechanism for the binding of off-target sequences at PAM distal sites, which has been unclear. Our simulations captured the interactions – formed between a locally unwound TS and the L2 loop – by which specific base pair mismatches prevent HNH activation for cleavage. This mechanism, which contributes in clarifying a long lasting open issue in the CRISPR-Cas9 function, also poses the basis for new rational engineering of the system toward improved genome editing. Indeed, structural modifications of the L2 loop – by using for instance non-natural amino acids – could be implemented with the goal of magnifying the “locking” interactions with the TS in the presence of off-target sequences. This could help the development of more specific Cas9 variants, in which a single base pair mismatch is sufficient for trapping HNH in its “conformational checkpoint”(12) state, thus preventing the cleavage of any incorrect DNA sequence.

## Method Summary

All simulations have been based on the X-ray structure of the *Streptococcus pyogenes* Cas9 in complex with RNA and DNA (4UN3.pdb), solved at 2.58 Å resolution.(5) A simulation protocol tailored for RNA/DNA endunucelases has been adopted,(26) embracing the use of the Amber ff12SB force field, which includes the ff99bsc0 corrections for DNA(30) and the ff99bsc0+χOL3 corrections for RNA.^(37, 38)^ To capture the conformational changes of the HNH domain in the presence of off-target sequences, an accelerated MD (aMD) method has been applied.(21) aMD is an enhanced sampling method that works by adding a non-negative boost potential to smoothen the system potential energy surface (PES), thus effectively decreasing the energy barriers and accelerating transitions between the low-energy states.(^21, 39^) Here, aMD simulations have been performed using the novel and more robust Gaussian aMD (or GaMD)(^23^) implementation, in which the boost potential follows Gaussian distribution, which extends the use of aMD to larger biological systems.(^24, 25, 40^) All simulations have been performed with the GPU version of AMBER 16.(41)

## Author contribution

G.P. designed this research and performed the simulations. G.P., C.G.R. and Y.M. analyzed the data. J.S.C and M.J. assisted in the interpretation of experimental data. J.A.M and J.A.D supervised research.

## Acknowledgments

G.P. thanks Alexis Komor for useful discussions. J.S.C thanks NSF for Graduate Research Fellowship. J.A.M. thanks NIH, NBCR and SDSC for support. This work used the Extreme Science and Engineering Discovery Environment (XSEDE), which is supported by National Science Foundation grant number ACI-1548562.42 This work used the XSEDE resource COMET at the San Diego Supercomputer Center through allocation TG- MCB160059.

